# *Symbiodinium* genomes reveal adaptive evolution of functions related to symbiosis

**DOI:** 10.1101/198762

**Authors:** Huanle Liu, Timothy G. Stephens, Raúl A. González-Pech, Victor H. Beltran, Bruno Lapeyre, Pim Bongaerts, Ira Cooke, David G. Bourne, Sylvain Forêt, David J. Miller, Madeleine J. H. van Oppen, Christian R. Voolstra, Mark A. Ragan, Cheong Xin Chan

**Affiliations:** Institute for Molecular Bioscience, The University of Queensland, Brisbane, QLD 4072, Australia; Australian Institute of Marine Science, Townsville, QLD 4810, Australia; ARC Centre of Excellence for Coral Reef Studies and Department of Molecular and Cell Biology, James Cook University, Townsville, QLD 4811, Australia; Laboratoire d’excellence CORAIL, Centre de Recherches Insulaires et Observatoire de l’Environnement, Moorea 98729, French Polynesia; Global Change Institute, The University of Queensland, Brisbane, QLD 4072, Australia; College of Science and Engineering, James Cook University, Townsville, QLD 4811, Australia; Research School of Biology, Australian National University, Canberra, ACT 2601, Australia; School of BioSciences, The University of Melbourne, VIC 3010, Australia; Red Sea Research Center, Division of Biological and Environmental Science and Engineering, King Abdullah University of Science and Technology (KAUST), Thuwal 23955-6900, Kingdom of Saudi Arabia; School of Chemistry and Molecular Biosciences, The University of Queensland, Brisbane,QLD 4072, Australia

## Abstract

Symbiosis between dinoflagellates of the genus *Symbiodinium* and reef-building corals forms the trophic foundation of the world’s coral reef ecosystems. Here we present the first draft genome of *Symbiodinium goreaui* (Clade C, type C1: 1.03 Gbp), one of the most ubiquitous endosymbionts associated with corals, and an improved draft genome of *Symbiodinium kawagutii* (Clade F, strain CS-156: 1.05 Gbp), previously sequenced as strain CCMP2468, to further elucidate genomic signatures of this symbiosis. Comparative analysis of four available *Symbiodinium* genomes against other dinoflagellate genomes led to the identification of 2460 nuclear gene families that show evidence of positive selection, including genes involved in photosynthesis, transmembrane ion transport, synthesis and modification of amino acids and glycoproteins, and stress response. Further, we identified extensive sets of genes for meiosis and response to light stress. These draft genomes provide a foundational resource for advancing our understanding *Symbiodinium* biology and the coral-algal symbiosis.

## Introduction

Coral reefs provide habitats for one-quarter to one-third of all marine species^1^. Although typically surrounded by nutrient-poor waters, coral reefs show high rates of primary productivity, with the fixed carbon supporting not only the biomass of reef organisms but also commercial and recreational fisheries and aquaculture. Reef-building corals rely on the symbiosis between the coral animal *per se* and photosynthetic dinoflagellates of the genus *Symbiodinium*. This symbiosis is based on mutual nutrient exploitation, with corals providing shelter and inorganic nutrients to their algal partners, while *Symbiodinium* supply their coral hosts with photosynthates that can meet up to 95% of the corals’ energy requirements^2,3^.

The relationship between *Symbiodinium* and their host determines not only the rate of coral-reef growth (calcium carbonate deposition), but also how the system responds to environmental stress^3^. Many studies have shown that coral-*Symbiodinium* mutualism is susceptible to environmental factors including temperature, light and salinity. Exposure to ultraviolet radiation, thermal stress or a combination of both can initiate photoinhibition, decoupling of carbon flow between symbiont and host, oxidative damage and breakdown of the symbiosis, a phenomenon known as coral bleaching^4^. Unless the symbiosis is soon re-established the coral host is at risk of starvation, disease and eventual death^5^. In recent decades, coral bleaching has led to large-scale mortality on coral reefs around the world, with the most recent global coral bleaching event (2014-2016) now confirmed as the longest and most severe on record^6,7^.

Despite the critical importance of this coral-dinoflagellate symbiosis, little is known about the underlying molecular mechanisms (apart from photosynthesis and carbon exchange), largely due to the lack of comprehensive understanding of what molecules, pathways and functions *Symbiodinium* can contribute. Genomes of *Symbiodinium* (and of dinoflagellates more broadly) are known for their idiosyncratic features including non-canonical splice sites, extensive methylation^8^ and large sizes, up to 250 Gbp^9^. Their plastid genomes occur as plasmid-like minicircles^10-12^; their mitochondrial genomes harbor only three protein-coding genes and lack stop codons, and both mitochondrial^13-15^ and nuclear^16^ transcripts are extensively edited.

*Symbiodinium* are classified into nine clades^17-19^, with members of Clades A, B, C and D responsible for the vast majority of associations with scleractinian corals^20^. Draft genomes have been published for representatives of Clades A, B and F^17-19^, with sequence comparisons demonstrating them to be highly divergent^18^. Genome sequences are still lacking for Clade C, the most ubiquitous and diverse clade associated with tropical reef corals^21^, at least some sub-clades (“types”) of which are ecologically partitioned^22^.

Here we report draft genomes of two *Symbiodinium* from the Pacific Ocean: *S. goreaui* (type C1; isolated from the acroporid coral *Acropora tenuis*) from the Great Barrier Reef, and *S. kawagutii* CS-156 (=CCMP2468, Clade F) from Hawaii. *Symbiodinium* type C1 is one of two “living ancestors” (along with type C3) of Clade C^21^, and one of the most dominant types associated with reef corals in both Indo-Pacific and Caribbean waters^20^. *S. goreaui* has been reported from >150 coral species on Australia’s Great Barrier Reef, representing > 80% of the studied coral genera in this region^23^ across environments from reef flats to lower mesophotic depths^23-25^. In contrast, *S. kawagutii* CS-156 (=CCMP2468) was isolated during attempts to culture the symbiont from *Montipora verrucosa* (Todd LaJeunesse, *personal communication*). This isolate has yet to be verified to occur in mutualistic symbiosis with any coral, and appears incapable of establishing experimental symbiosis with cnidarian hosts^26^. Instead *S. kawagutii* may be exclusively a symbiont of foraminifera, or occur free-living at low environmental densities but proliferate opportunistically in culture. As some genome data have been published for *S. kawagutii* CCMP2468^18^, we used these in combination with new data from the present study to generate a refined genome assembly. The genomes of *S. goreaui* and *S. kawagutii* offer a platform for comparative genomic analyses between two of the most-recently diverged *Symbiodinium* lineages Clades C and F, and published genome sequences in the more-basal Clades A and B.

Adopting a comparative approach using both genome and transcriptome data, we systematically investigated genes and functions that are specific to *Symbiodinium* vis-à-vis other dinoflagellates, and their association with the establishment and maintenance of symbiosis. We also computationally identify genes and functions for which there is evidence of adaptive selection in *Symbiodinium*. This is the most-comprehensive comparative analysis so far of *Symbiodinium* genomes, and the first to include a prominent endosymbiont of corals of Indo-Pacific and Caribbean reefs.

## Results

### Genomes of S. goreaui and S. kawagutii

We sequenced and generated two draft *Symbiodinium* genome assemblies *de novo*, for *S. goreaui* (Clade C, 1.03 Gbp) and for *S. kawagutii* (Clade F, 1.05 Gbp). Details of data generation and assembly statistics are shown in Supplementary Tables S1 and S2 respectively. Our *S. goreaui* assembly consists of 41,289 scaffolds (N50 length 98,034 bp). For *S. kawagutii*, we first verified that our data (from isolate CS-156) and the published data (from the synonym isolate CCMP2468) are indeed from the same culture of origin (see Supplementary Methods and Supplementary Figure S1). Compared to the published assembly (Lin et al.^18^), independent mapping of their ten fosmid sequences^18^ onto our preliminary CS-156 assembly yielded up to 43-fold and 37-fold fewer gaps and mismatches respectively (Supplementary Figure S2). We later combined both datasets in a single *de novo* assembly,yielding 16,959 scaffolds (N50 length 268,823 bp). Genome-size estimates based on *k*-mer coverage are 1.19 Gbp for *S. goreaui* and 1.07 Gbp for *S. kawagutii* (Supplementary Table S3), comparable to those for other sequenced *Symbiodinium* genomes. We also recovered sequences putatively derived from their plastid genomes (Supplementary Tables S4, S5 and S6) including their distinct core conserved regions (Supplementary Table S7), and from their mitochondrial genomes; see Supplementary Note for details.

The repeat content of the assembled genomes ranged from 16.0% (*S. kawagutii*) to 27.9% (*S. microadriaticum*); a large peak in transposable element (TE) abundance observed at high divergence (Kimura distance^27^ 15-25) in all genomes (Supplementary Figure S3) suggests that most extant transposable elements are remnants of an ancient burst of TE activity. TE activity has been broadly linked to genome size in plants^28-30^, so reduced TE activity may be connected with the relative compactness of *Symbiodinium* genomes in comparison to those of other dinoflagellates. However, as these genomes are still in draft, the impact of assembly completeness on the patterns of repeat divergence cannot be dismissed.

Using a stringent threshold to remove genome scaffolds of potential bacterial or viral origin (Methods), we predict 35,913 and 26,609 high-quality gene models respectively for *S. goreaui* and *S. kawagutii* (Supplementary Table S8). Usage profiles of codons and amino acids are shown in Supplementary Figures S4 and S5 respectively, and non-canonical splice sites in Supplementary Table S9 and Supplementary Figure S6. Although we report fewer genes, the majority (67.0% and 64.4% respectively for *S. goreaui* and *S. kawagutii*) have transcriptome support, and we generally recovered more of the 458 conserved core eukaryote genes (Supplementary Figure S7), 371 of which are common to all four *Symbiodinium* based on the predicted gene models (Figure 1a; Supplementary Table S10); similar results are observed for the corresponding genome sequences (Supplementary Figure S7). About 94% of the predicted genes have introns, similar to *S. microadriaticum* (98.2%) and *S. minutum* (95.3%); the earlier *S. kawagutii* genome assembly^18^ may have underestimated the proportion of intron-containing genes (Supplementary Table S8) due to a less-stringent approach to gene prediction. All coding sequences have higher G+C content (56.7% in *S. goreaui* and 55.0% in *S. kawagutii*) than does the genome overall, comparable to coding sequences from other *Symbiodinium* (57.7% in *S. microadriaticum* and 52.7% in *S. minutum*).

**Figure 1.**
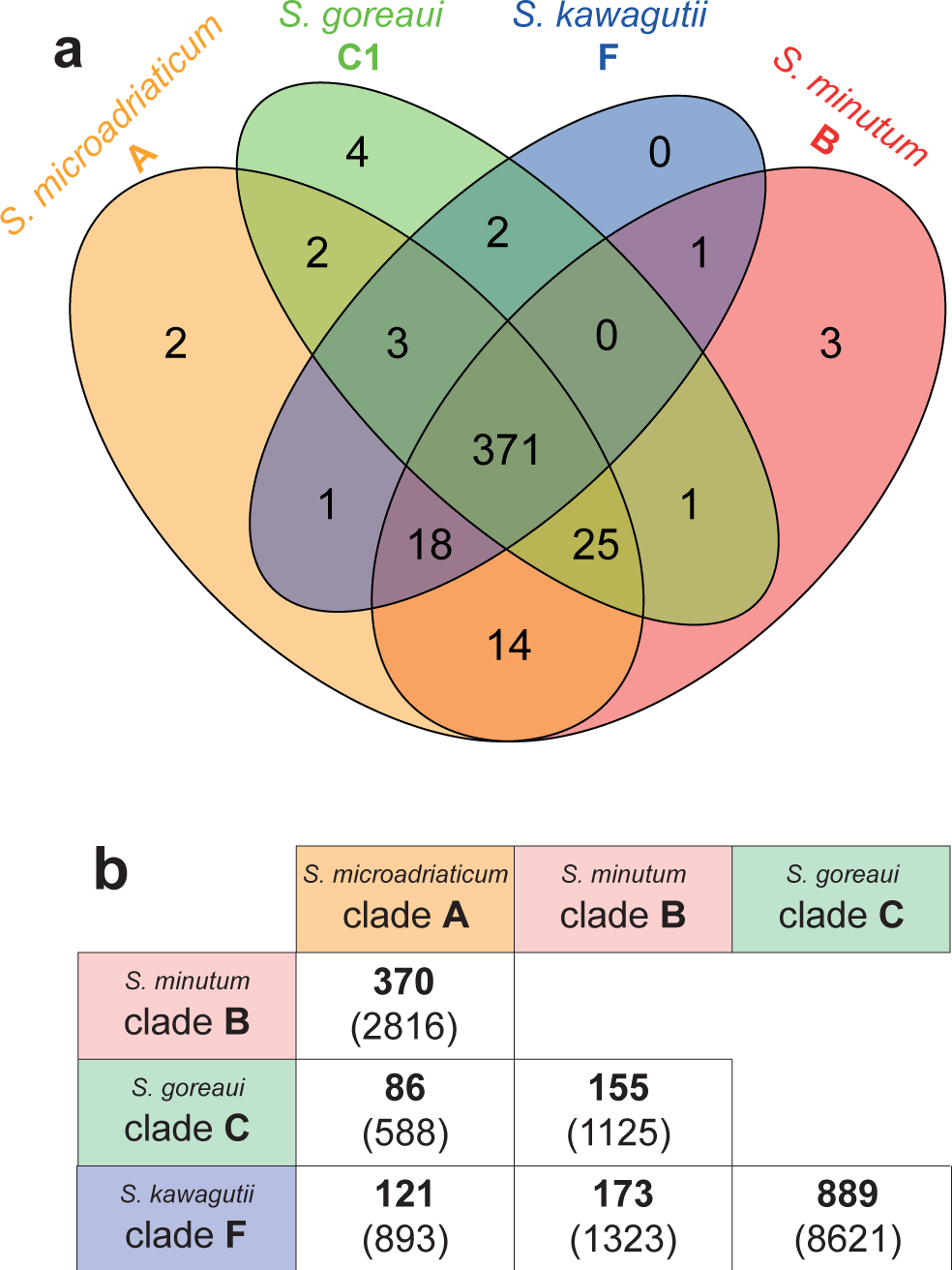
Comparison of *Symbiodinium* genomes. (**a**) Number of recovered core eukaryote genes in each genome based on CEGMA. (**b**) Number of identified syntenic collinear blocks (and the corresponding number of implicated genes) between each genome pair.

### Sequence divergence and synteny

Despite the seemingly high number of protein-coding genes, an earlier analysis of syntenic blocks^17^ found only several hundred blocks conserved in a pairwise manner among three published *Symbiodinium* genomes. Here we included our two new genome sequences in this analysis, and focused further on syntenic collinear blocks, requiring each block to contain the same genes in the same order and orientation with no gene losses (Methods). The *S. goreaui* and *S. kawagutii* genomes share the most collinear blocks, 889 blocks implicating 8621 genes; 62 of these blocks are of size >15, with the largest containing 76 genes (Supplementary Table S11). Thus substantial proportions of genes in these two genomes occur in clusters: for cluster size ≥ 5 genes, 32.4% and 24.0% of *S. kawagutii* and *S. goreaui* genes respectively. These are likely to be underestimates, as the assemblies remain fragmentary. At the other end of the spectrum, the genomes of *S. microadriaticum* and *S. goreaui* share only 86 collinear blocks of size ≥ 5, with maximum size 12 and implicating 588 genes in total (Supplementary Table S11). These results suggest that (a) although Clades C and F are divergent, they nonetheless are the most-closely related among the four analysed *Symbiodinium* genomes (in line with their phylogenetic relationship); and (b) C and F are more divergent from Clade A than from Clade B (in line with their phylogenetic relationship). Thus whole-genome sequences support and extend earlier conclusions, based on common marker sequences, about phylogenetic relationships among *Symbiodinium* clades^31-33^ and the remarkable divergence among *Symbiodinium* lineages^17,18^.

### Gene and protein functions

All annotated genes from *S. goreaui* and *S. kawagutii* genomes are listed in Supplementary Tables S12 and S13 respectively. Of the 35,913 proteins predicted in *S. goreaui*, 31,646 (88.1%) show similarity (BLASTP, *E* ≤ 10^−5^) to sequences in UniProt; among these, 29,198 (81.3% of 35,913) and 19,718 (54.9%) are annotated with Gene Ontology (GO) terms or Pfam domains (Supplementary Tables S12 and S14). For *S. kawagutii*, 21,947 of 26,609 proteins (82.5%) find a match in UniProt (Supplementary Tables S13 and S14). *Protein kinase* (Pfam PF00069), *reverse transcriptase* (PF07727), *ion transport protein* (PF00520) and *ankyrin repeats* (PF12796) are among the most-abundant domains in *Symbiodinium*, appearing among the ten most-abundant for each of the four genomes (Supplementary Table S15). Ankyrin repeat motifs are important in the recognition of surface proteins, and more generally in protein-protein interactions^34^. Thus proteins potentially involved in host-symbiont interaction (with phosphorylation, ion transport and protein recognition/interaction domains) are well represented in the predicted *Symbiodinium* proteomes.

We compared functions of proteins predicted from these four *Symbiodinium* genomes to a set of 27 more-narrowly scoped eukaryotes: 17 alveolates (ten other dinoflagellates, four ciliates, two apicomplexans and *Perkinsus marinus*), stramenopiles (two diatoms) and Archaeplastida (four rhodophytes, three chlorophytes and *Arabidopsis*). This 31-taxon set (1,136,347 proteins; Supplementary Tables S16 and S17) represents lineages in which one or more endosymbioses are implicated in plastid origin^35-37^; these proteins were clustered (based on sequence similarity) into 56,530 groups of size two or greater (Supplementary Table S17; see Methods). Using this 31-taxon dataset as background, we assessed the over- or under representation of protein domains within our various groups of *Symbiodinium* proteins. We found 270 domains (Supplementary Table S18) to be significantly overrepresented (adjusted *p* ≤ 0.05) in *Symbiodinium*. Interestingly, many domains e.g. *C-5 cytosine-specific DNA methylase* (PF00145), *planctomycete cytochrome c* (PF07635) and RNA polymerase *sigma-70 region 2* (PF04542) are also overrepresented in the four *Symbiodinium* genomes in a similar comparison against 880,909 proteins in a 15-taxon set that includes ten other dinoflagellates and the immediate outgroup *Perkinsus marinus* (Supplementary Table S19). Thus compared to related eukaryotes and to other dinoflagellates, *Symbiodinium* is enriched in functions involved in methylation of cytosine, (photosynthetic) energy production and RNA polymerisation. Hydroxymethylation of uracil is common (12-70%) in dinoflagellate genomes^38^; while cytosine methylation has been described in *Symbiodinium*^39,40^, our findings suggest that cytosine methylation is more prominent in *Symbiodinium* than in these other dinoflagellates.

Activation of some retrotransposons is part of the stress-response mechanism in diatoms^41^, plants^42,43^ and other eukaryotes^44,45^. The *reverse transcriptase* domain (PF07727) is enriched in *Symbiodinium* in both the 31-taxon and 15-taxon sets, suggesting that retrotransposition could be a prominent mechanism of stress response in *Symbiodinium* and dinoflagellates. Although we set a stringent threshold for removing putative bacterial or viral sequences (see Methods), 40 (∼0.1%) of the final 41,289 genome scaffolds of *S. goreaui* have significant hits (BLASTN *E* ≤ 10^−20^) to the virus genomes^46^ isolated from the same *S. goreaui* (type C1) strain, with 16 identical regions (76-609 bp) distributed in nine scaffolds of lengths ranging from 1092 to 7,338,656 bp. Whether this indicates introgression of viral sequences remains to be determined.

### Positive selection of Symbiodinium genes

Using a branch-site model based on the ratio of substitution rates in non-synonymous (dN) to synonymous (dS) sites^47^ (Methods and Supplementary Figure S8), we identified *Symbiodinium* genes under positive selection in comparison to ten other dinoflagellates, with *Perkinsus marinus* as the outgroup (15 taxa: Supplementary Tables S16 and S17). The reference species tree (Figure 2a) was computed following Price and Bhattacharya^48^. We then based our analysis of adaptive evolution on all orthologous sets for which the protein tree is topologically congruent with our reference tree (Methods).

**Figure 2.**
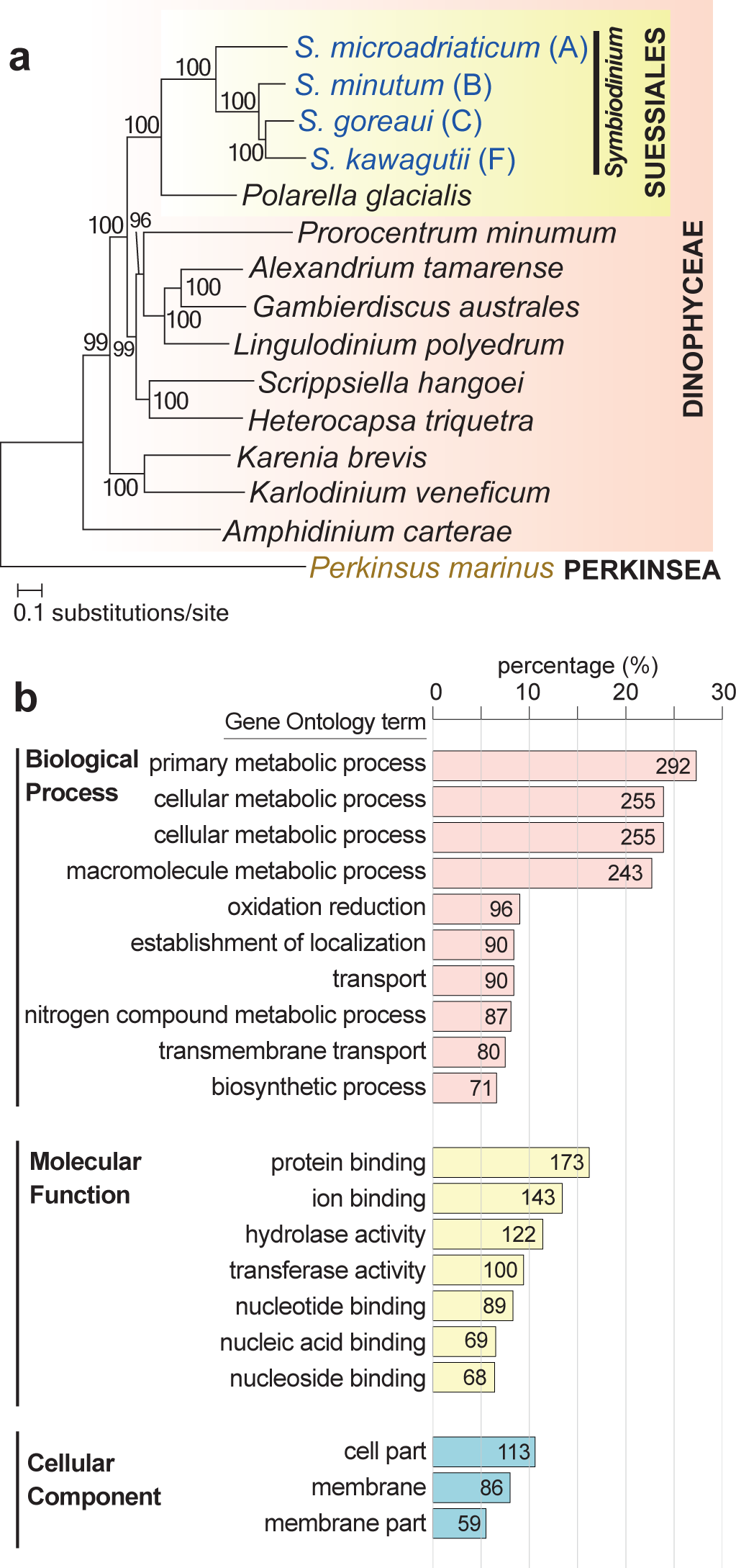
Testing for positive selection acting on *Symbiodinium* genomes. (**a**) The reference 15-species supertree of *Symbiodinium*, dinoflagellates and *Perkinsus marinus* (as outgroup) based on single-copy orthologous genes. Support based on 2000 rapid bootstraps is shown on each internal node, and the branch length is the number of substitutions per site. (**b**) Percentage of positively selected genes with annotated GO (level 3) terms in *Symbiodinium*, shown for principal hierarchies Biological Function, Molecular Function and Cellular Component.

Of our 44,282 homologous sets, 2460 containing 7987 *Symbiodinium* proteins show evidence of positive selection in one or more *Symbiodinium* lineages; 1069 of these sets are annotated with GO terms (Supplementary Table S20). Figure 2b shows the terms (level 3) in the three GO hierarchies that are shared by ≥ 5% of these 1069 sets. In the Biological Process hierarchy, metabolic processes are highly represented (*primary metabolic process* [292] and *macromolecule compound metabolic process* [243] are among the four most-frequent terms), followed by *oxidation reduction* [96] and transport (*establishment of localization* [90], *transport* [90], and *transmembrane transport* [80]). Highly represented terms in the Molecular Function hierarchy point to binding of diverse molecules and ions e.g. *protein binding* [173] and metabolism (*hydrolase* [390], *transferase* [344]). In Cellular Component, *cell part* [113], *membrane* [86] and *membrane part* [59] are the most frequent. Thus in *Symbiodinium* as represented by these four assemblies, broad aspects of metabolism, and transport including across membranes, show evidence of positive selection, in line with their recognised importance in cnidarian-dinoflagellate symbioses^17^.

We further assessed enrichment of GO terms against all annotated terms specifically in the four *Symbiodinium* genomes (Supplementary Table S21) compared with the other dinoflagellates in this study. Here we consider themes among Biological Process terms. The first theme is that functions associated with photosynthetic light reactions are enriched among the positively selected *Symbiodinium* genes; *photosynthesis, light reaction* and *Photosystem II assembly* are significantly over-represented (adjusted *p* ≤ 0.05), as are Cellular Component terms related to plastid functions e.g. *thylakoid, photosynthetic membrane, intracellular membrane-bounded organelle* (Supplementary Table S21). Coral bleaching has been associated with the loss of light-harvesting proteins and the subsequent inactivation of photosystem II (PSII) in *Symbiodinium* under combined light and temperature stress^49-51^.These authors reported that coral bleaching associated with algal photobleaching can be ameliorated, at least in part, by thermal acclimation of *Symbiodinium* to improve the thermal tolerance of PSII.

The second emerging theme involves the transport of ions and metabolites across membranes. *Intracellular transport, cytosolic transport, transition metal ion transport* and *copper ion transport* as well as terms related to transmembrane transport of amino acids, organic acids and carboxylic acids are significantly enriched (Supplementary Table S21); these functions underpin multiple physiological processes, including but not limited to pH regulation, calcification and photosynthetic carbon fixation^52^. Investigated *Symbiodinium* are enriched in membrane transporters compared with other dinoflagellates^17^. These biological processes are especially relevant to the maintenance and regulation of coral-dinoflagellate symbiosis^52,53^, possibly including its sensitivity and/or response to environmental stress.

The third theme is the enrichment of functions related to the biosynthesis and modification of amino acids and glycoproteins (Supplementary Table S21) e.g. *protein phosphorylation, peptide biosynthesis process, protein ADP-ribosylation, protein glycosylation, D-amino acid metabolic process* and *glycoprotein biosynthetic process*. Corals lack the capacity to synthesise a number of amino acids (e.g. cysteine in *Acropora digitifera*^54^), thus selection acting on the synthesis of amino acids may indicate the critical role of *Symbiodinium* in supplying amino acids both for self-preservation and for the coral host. Glycoprotein molecules are often surface-localised and in microbes form the basis of microbe-associated molecular patterns (MAMPs) which, in conjunction with a host-associated pattern recognition receptor, mediate recognition by a host^55^. Both in culture and *in hospite, Symbiodinium* exude glycoconjugates^56-59^. Where investigated, cell-surface glycan profiles are stable over time within a *Symbiodinium* culture but can differ between clades within a host^60^. *N*-acetyl and mannosyl residues are prominent constituents of *Symbiodinium* cell-surface glycans, making them candidates for MAMPs that could participate in the establishment of symbiosis. Lin et al.^18^ reported a *S. kawagutii* glycan biosynthesis pathway distinct from that of *S. minutum*, again pointing to a possible role of glycans in specificity of host recognition^60,61^. Neubauer et al.^62^ demonstrated that the thrombospondin type 1 repeat (TSR) from the sea anemone *Aiptasia pallida* contains binding sites for glycosaminoglycan, and that blocking TSR led to decreased colonisation by *S. minutum*. Our results offer the first evidence of positive selection of functions underlying the biosynthesis and modification of amino acids and glycoproteins, suggesting that these functions are critical in the establishment of cnidarian-dinoflagellate symbioses.

Our fourth emerging theme is stress response. Enriched terms annotated for the positively selected genes include *cell redox homeostasis, translation initiation* and 22 terms describing the negative regulation of gene expression, transcription, RNA biosynthesis and cellular biosynthetic and metabolic processes (Supplementary Table S21). Environmental stressors can disrupt the cellular redox homeostasis and break down the coral-dinoflagellate symbiosis. Negative regulation of transcription may represent a stress response that safeguards the genome from accumulating DNA damage^63^; a similar stress response has been observed in coral^64^. Other enriched functions that may be related to stress response include *mitotic nuclear division, translation*, and various processes of nucleotide biosynthesis and modification e.g. *RNA methylation, rRNA methylation, DNA replication, RNA processing,* and *deoxyribonucleotide biosynthetic process*. Our results thus provide evidence that stress response is under positive selection in *Symbiodinium*, presumably (given their lifestyle) in relation to the establishment and/or maintenance of symbiosis.

### Do Symbiodinium have sex?

*Symbiodinium* have been hypothesised to reproduce sexually and to have a diploid life stage^65^ but definitive evidence for sex, e.g. karyogamy and meiosis, has yet to be observed^31,66-68^.The ability to reproduce sexually offers increased efficiency of selection and adaptation^69^. So far, the strongest evidence supporting meiotic potential in *Symbiodinium* comes from patterns of population-genetic variation revealed in allozymes, randomly amplified polymorphic DNA and other molecular markers^22,31,70-73^. Indeed, for some markers a higher degree of genetic variation has been observed in certain *Symbiodinium* clades than in dinoflagellates known to reproduce sexually^70^. More recently, differential gene expression analysis^74^ using a heterologous culture from which our sequenced *S. goreaui* was derived revealed an enrichment of gene functions related to meiosis under thermal stress, suggesting a switch from mitosis to meiosis under stress conditions.

Schurko and Logsdon^75^ described a “meiosis detection toolkit”, a set of 51 genes specific or related to meiosis^76,77^ that collectively point to a capacity for meiosis even in divergent or specialised eukaryotic genomes^78^. Incomplete genome coverage or assembly, sequence divergence, paralogy, patterns of overlapping function and evolutionary specialisation mean that not all 51 need to be present or detectable for a lineage to be assessed as probably sexual, or only recently asexual^75,77^. Thirty-one of these genes were earlier identified in *Symbiodinium* Clades A and B^76^. Here, BLASTP search (*E* ≤10^−5^) of predicted proteins in these four *Symbiodinium* genomes recovered matches corresponding to 48 of the of 51 toolkit genes in *S. microadriaticum*, 47 in *S. minutum* and *S. goreaui*, and 46 in *S. kawagutii* (Figure 3a and Supplementary Table S22). Eight of the 11 meiosis-specific proteins were detected in all four *Symbiodinium*. REC114, SAD1 and XRS2 found weaker matches (*E* ≥ 10^−14^) in one to two genomes, although confirmatory UniProt domains were usually present (Supplementary Table S22). RAD17 is the *Schizosaccharomyces pombe* homolog of *S. cerevisiae* RAD24^79^, for which we find highly significant matches (*E* ≤ 10^−127^) in all four *Symbiodinium*. Moreover, 15 of the 51 genes show evidence of positive selection in *Symbiodinium* against other dinoflagellates (Supplementary Table S22). Our data imply that these four *Symbiodinium* are, or until recently have been, capable of meiosis.

**Figure 3.**
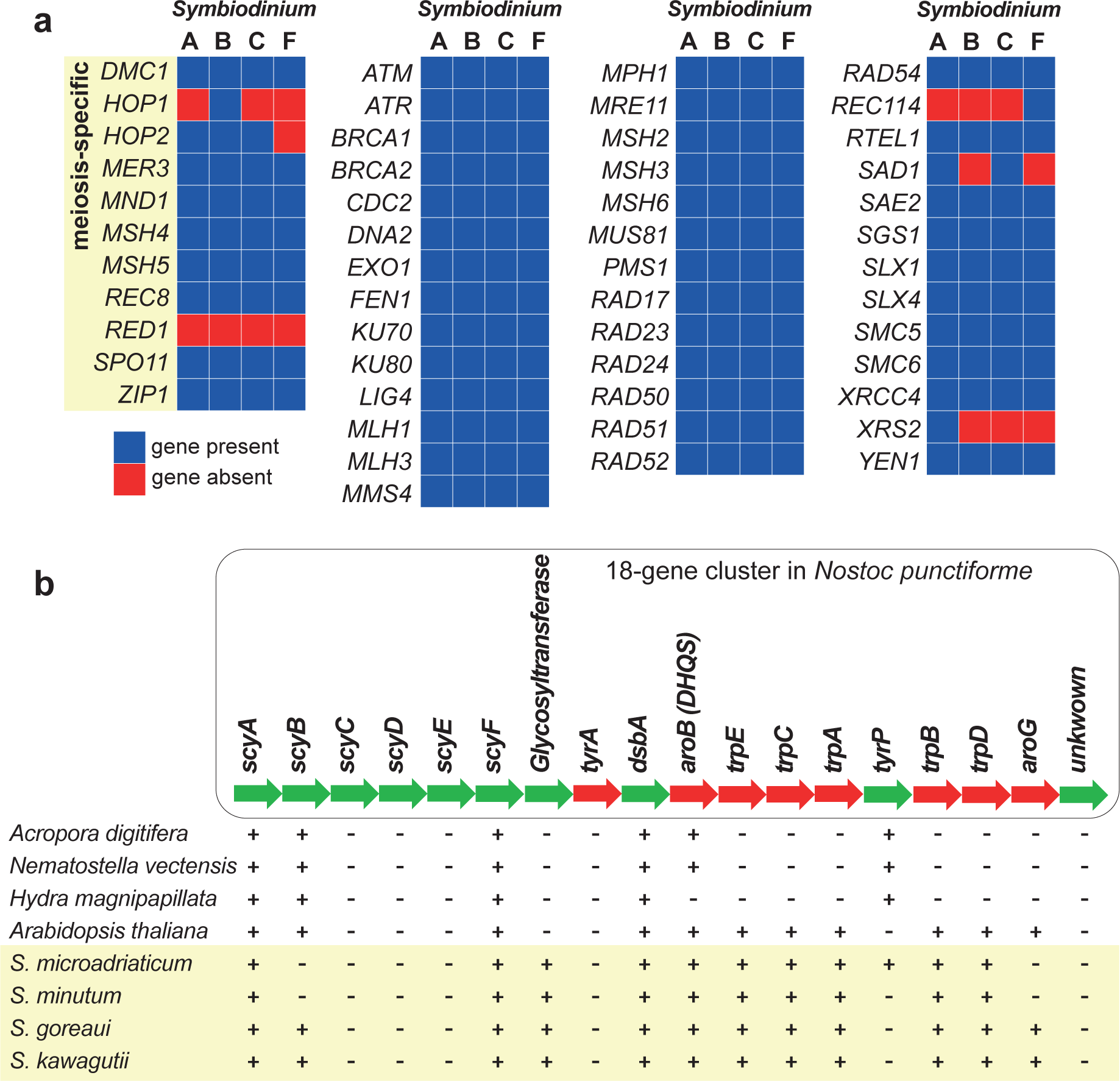
Recovery of genes in *Symbiodinium*. (**a**) Meiosis-related genes recovered in the genomes of *S. microadriaticum* (Clade A), *S. minutum* (Clade B), *S. goreaui* (Clade C) and *S. kawagutii* (Clade F). (**b**) Scytonemin biosynthesis genes in *Symbiodinium* genomes relative to the coral *Acropora digitifera*, sea anemone *Nematostella vectensis*, hydra (*Hydra magnipapillata*) and the green plant *Arabidopsis thaliana*. The order of the 18-gene cluster in the cyanobacteria *Nostoc punctiforme* is used as a reference, with presence (+) and absence (-) of a gene in each species are shown. Figure modified from Shinzato et al.^90^.

### Response to light stress

Mycosporine-like amino acids (MAAs) are UV-protective compounds that, in corals and other marine organisms, also act as antioxidants scavenging reactive oxygen species (ROS). Up to five MAAs have been reported in *Symbiodinium* (Clades A, B and C) isolated from cnidarian hosts^80,81^. The MAA biosynthetic pathway involves dehydroquinate synthase (DHQS), *O*-methyltransferase (O-MT), an ATP-grasp and non-ribosomal peptide synthetase (NRPS)^82,83^. In cyanobacteria, a short-chain dehydrogenase may play a role in converting shinorine to palythine-serine^84^. Genes encoding these four MAA-biosynthetic enzymes were reported absent from the *S. kawagutii* genome^18^. Here, using known proteins in bacteria, fungi and cnidarians as queries, we recovered all five enzymes including the short-chain dehydrogenase from the *S. microadriaticum, S. goreaui* and *S. kawagutii* genomes (Supplementary Table S23); ATP-grasp was not found in *S. minutum*. These enzymes were earlier reported absent from *S. kawagutii*, and it was proposed that their absence can be compensated via coral-*Symbiodinium* co-evolution^18^; this hypothesis remains to be investigated, but we note that this *S. kawagutii* isolate has not been observed in association with an animal host^26^.

Scytonemin is a UV-blocker first reported in terrestrial cyanobacteria^83,85^, and in contrast to MAAs was thought to be synthesised exclusively by cyanobacteria^86^. The genome of *Nostoc punctiforme* contains an 18-gene operon that specifies proteins of scytonemin biosynthesis and regulation, including the synthesis of aromatic amino-acid precursors^87,88^. Its expression is up-regulated by UV radiation^89^. Homologs of six of these 18 genes have been described in the coral *Acropora ditigifera*, and were considered putative instances of lateral genetic transfer^90^. We find 12 of these 18 genes in the genomes of *S. goreaui* and in *S. kawagutii*, 11 in *S. microadriaticum* and ten in *S. minutum* (Figure 3b, Supplementary Table S24).

Genes responsible for biosynthesis of tryptophan (*trpA, trpB, trpC, trpD* and *trpE*) and the two key enzymes of chorismate (and aromatic amino acid) biosynthesis, *aroG* and *aroB* (dehydroquinate synthase, also important for MAA biosynthesis), are found in all *Symbiodinium* genomes, albeit so far in different scaffolds; these genes are also present in *Arabidopsis thaliana* although not in corals or *Hydra* which, like most other animals, are unable to synthesise tryptophan. The recovery of more of these 18 genes in *Symbiodinium* than in corals or other animals (Figure 3b) could reflect the impact of endosymbiotic association of ancestral cyanobacteria during the course of plastid evolution in all photosynthetic eukaryotes including dinoflagellates^35-37^. The presence of multiple gene copies (Supplementary Table S24) also implicates genetic duplication. Our findings suggest that *Symbiodinium* has the capacity to produce scytonemin, either by itself or jointly within the holobiont.

## Discussion

*Symbiodinium* can form associations with a wide range of cnidarian hosts (as well as some other marine invertebrates and protists) across broad geographic and time scales^91^. The symbiosis between corals and *Symbiodinium* relies on compatible host-symbiont recognition and sustainable nutrient exchange, both of which are vulnerable to external environmental factors including temperature and light. A sustainable coral-*Symbiodinium* association requires an adaptive capacity in the face of environmental extremes.

In this study we generated the first draft genome of S. goreaui (Clade C), a much-improved draft genome of S. kawagutii (Clade F) and high-quality gene models for both. Comparative analysis revealed remarkable divergence among the genomes of Symbiodinium from four clades, consistent with previous single-gene phylogenetic relationships. We identified 2460 Symbiodinium gene families that are under positive selection, many of which encode functions directly relevant to the establishment and/or maintenance of symbiosis with the coral host. We also identified complete, or near-complete, sets of genes indicative of the presence of meiosis and several mechanisms of stress tolerance, functions that have until now been considered absent from S. kawagutii. Our results demonstrate the remarkable genomic capacity of Symbiodinium to synthesize key metabolites that are essential to the establishment of symbiosis with coral hosts, and to respond to environmental stress.

*S. goreaui* (type C1) belongs to one of the most globally dominant clades (Clade C) on coral reefs, and analysis of its draft genome has provided important new insights into coral-algal symbiosis. This genomic resource is already demonstrating utility in the identification of symbiont fractions in *de novo* sequencing of coral tissues^92,93^, and will provide a foundation for targeted studies into the molecular biology, physiology and of this crucial symbiosis and its adaptation to a changing environment.

## Methods

### Biological materials and sequencing data

*Symbiodinium goreaui* (Clade C, type C1; AIMS-aten-C1-MI-cfu-B2, now AIMS culture collection SCF055-01) is a single-cell monoclonal culture first isolated from the coral *Acropora tenuis* at Magnetic Island (Queensland, Australia) at 3 m depth^94^. *Symbiodinium kawagutii* CCMP2468 (Clade F) was maintained as a monoclonal culture. Genomic DNA was extracted from these isolates using the Qiagen DNeasy Plant Mini Kit. We generated a total of 116.0 Gb of sequence data (2 paired-end libraries of 230-and 500-bp inserts, plus 3 mate-pair libraries of 3-,6- and 9-Kbp inserts) for *S. goreaui*, and a total of 92.2 Gb of sequence data (1 paired-end library of 230-bp inserts, plus 3 mate-pair libraries of 4-, 6-and 9-Kbp inserts) for *S. kawagutii*, in each case using the Illumina HiSeq 2500 Rapid Chemistry platform. See Supplementary Methods and Supplementary Table S1 for details.

### Genome assembly and annotation

We adopted a combined approach to *de novo* genome assembly (Supplementary Methods) in which multiple assembly programs (CLC Genomics Workbench (Qiagen), SPAdes^95^ and ALLPATHS-LG^96^) were first used independently. The quality of each assembly was assessed based on full-length recovery of phylogenetic markers and known coding sequences. Once the best assembly (the master assembly) was identified, other assemblies were used to refine it via merging scaffolds and filling gaps. We adopted a comprehensive *ab initio* approach for gene prediction using all available dinoflagellate proteins, as well as all *Symbiodinium* genes and transcriptomes, as guiding evidence. Our approach combines evidence-based methods i.e. PASA (with TransDecoder)^97^, AUGUSTUS^98^ and MAKER^99^, and unsupervised machine-learning (GeneMark-ES^100^ and SNAP^101^; alternative splice sites were specified in these methods by modifying the relevant scripts. We then used EvidenceModeler^102^ to combine multiple sets of predicted genes, allocating a heavier weight for evidence-based predictions than for those produced by unsupervised approaches; for details see Supplementary Methods. Final genome assemblies, predicted gene models and proteins are available at https://cloudstor.aarnet.edu.au/plus/index.php/s/6yziMf2ygWjGu0L.

We adopted multiple approaches to assess genome completeness. Established methods including CEGMA^103^ and BUSCO^104^ are based on conserved genes in a limited number of eukaryote model organisms that are distantly related to dinoflagellates. The use of these methods resulted in relatively low recovery of conserved eukaryote genes in *Symbiodinium* (e.g. 33-42% of BUSCO genes; Supplementary Figure S7) when run at default setting. We further assessed completeness using BLAST based on predicted proteins from the gene models and the assembled genome scaffolds. For each genome, we followed Baumgarten et al.^105^ and searched (BLASTP, *E* ≤10^−5^) against the predicted proteins using the 458 CEGMA proteins^103^. We also searched against the CEGMA proteins using the genome scaffolds (BLASTX *E* ≤ 10^−5^), against genome scaffolds using the 458 CEGMA proteins (TBLASTN, *E* ≤ 10^−5^), and against genome scaffolds using the 458 CEGMA transcripts (TBLASTX, *E* ≤ 10^−5^) (Supplementary Table S10, Supplementary Figure S7).

### Identification and removal of bacterial and viral sequences

Bacterial and viral sequences were identified and removed following Aranda et al.^17^. Briefly, we used our genome scaffolds (BLASTN, *E* ≤ 10^−20^) to query the complete and draft bacterial genomes in NCBI, and the viral genomes in NCBI and PhAnToMe (http://phantome.org). We applied more-stringent criteria than did Aranda et al.^17^ to identify putative contaminating sequences, removing from the final assembly any scaffold sequence that contains regions that match a bacterial or viral genome (BLASTN bit score > 1000, *E* ≤ 10^−20^) where such regions constitute >10% of the overall scaffold length (Supplementary Methods, Supplementary Figure S9).

### Analysis of genome synteny and collinearity

Using all predicted genes and their associated genomic positions, we assessed the number of syntenic collinear blocks (i.e. regions with the same genes coded in the same order, free from rearrangement or loss) shared pairwise among genomes of *S. microadriaticum* (Clade A)^17^, *S. minutum* (B)^19^, *S. goreaui* (C) and *S. kawagutii* (F). We used BLASTP (*E* ≤ 10^−5^) to search for similar proteins within each genome, and among all of them. Next we used MCScanX^106^ with parameter -s 5 to sort the BLASTP matches (alignments) based on genomic positions; to minimise the number of collinear gene pairs arising from tandem repeats, we report only collinear blocks that consist of five or more genes.

### Functional annotation of gene models

We adopted a similar approach to Aranda et al.^17^ to annotate gene models. Protein sequences predicted using the standard genetic code were used as query (BLASTP, *E* ≤ 10^−5^) first against Swiss-Prot, and those with no Swiss-Prot hits subsequently against TrEMBL (both databases from UniProt release 2016_10). Gene Ontology (http://geneontology.org/) terms associated with Swiss-Prot and TrEMBL hits were obtained using the UniProt-GOA mapping (release 2016_10).

### Identification of homologous protein groups and gene families

Our workflow for delineation of sets of putatively homologous proteins, multiple sequence alignment, generation of protein-family and reference trees, and analysis of selection is shown in Supplementary Figure S8. Using predicted proteins from 31 phyletically diverse organisms including *Symbiodinium* for which genome and/or transcriptome data are available (Supplementary Table S25), we identified sets of putatively homologous proteins using OrthoFinder^107^ and consider the corresponding gene sets (families) to be homologs. Following Harlow et al.^108^ and Beiko et al.^109^, we consider sequences within single-copy sets (i.e. those in which each genome is represented no more than once) to be orthologs. We refer to sets that contain proteins only from *Symbiodinium*, plus the *Symbiodinium* singletons, as *Symbiodinium*-specific. For enrichment analysis of annotated features (GO terms or Pfam domains), we compared the features within the *Symbiodinium*-specific set against those in each background set (i.e. the 31-taxon set and, separately, the 15-taxon set) using a hypergeometric test; features with an adjusted^110^ *p*-value < 0.05 were considered significant.

### Analysis of positive selection in Symbiodinium genes

For the 15-taxon set we sorted the 311,651 protein sets into 1,654 single-copy (ortholog) and 16,836 multi-copy sets. Multiple sequence alignments were carried out using MAFFT v7.245 at -linsi mode^111^; questionably aligned columns and rows were removed from these alignments using trimAl^112^ with the -automated1 option.

Branch-site models (BSMs; see below) require a reference topology. We follow Price and Bhattacharya^48^ to generate the reference species tree. The trimmed single-copy protein alignments were concatenated prior to maximum-likelihood (ML) inference of the species phylogeny using IQTREE^113^; each alignment represents a partition for which the best evolutionary model was determined independently. Support for each node was assessed using 2000 rapid bootstraps. The species tree so generated (Figure 2a) is similar to that of Price and Bhattacharya^48^, with very strong support (≥ 96% bootstrap replicates) for all internal nodes; the *Symbiodinium* and Suessiales (*Symbiodinium* + *Polarella glacialis*) clades received 100% bootstrap support.

Of all trimmed protein alignments, those with ≥ 60 aligned positions and ≥ 4 sequences were used in subsequent analysis. For multi-copy protein sets, we imposed further filtering criteria. We first inferred individual ML trees for the multi-copy sets using IQ-TREE, and each resulting protein tree was compared with the reference species tree. Those congruent with the reference species tree at genus level, and in which all *Symbiodinium* are resolved as an exclusive monophyletic clade, were judged paralog-free and used in subsequent BSM analysis (Supplementary Figure S8). Among the 16,836 multi-copy sets of the 15-taxon analysis, 1788 (10.6%) resolve all *Symbiodinium* sequences into an exclusive monophyletic clade and are topologically congruent at genus level with the reference species tree (i.e. contain co-orthologs but not paralogs) and were retained, while the remaining 15,048 failed one or both of these filtering criteria (i.e. contain presumed paralogs) and were not analysed further (Supplementary Figure S8). The percentages of missing data and parsimoniously informative sites in all filtered protein alignments for the 15-taxon set are detailed in Supplementary Table S26. For each filtered alignment, we used the corresponding coding-sequence alignment (codon alignment) generated using PAL2NAL^114^ in the BSM analysis. Some predicted protein sequences in MMETSP^115^ do not match their corresponding CDS, sometimes due to problematic translation and other times due to a frameshift. For these, we used MACSE^116^ to derive the codon alignments.

We applied the branch-site model (BSM) implemented in the *codeml* program in PAML 4.9^117^ to detect positive selection signal unique to the *Symbiodinium* lineage. BSMs allow the dN/dS ratio ω to vary among both sites and branches, making it possible to infer selection at both. We computed two models: a null model with fixed ω = 1, and an alternative model that estimates ω in our defined foreground branches (here, the node that leads to all *Symbiodinium* lineages). We then compared the likelihoods of these two models to determine the better fit. To reduce false positives we applied *q*-value estimation for false discovery rate control, as implemented in R package *qvalue* to adjust *p*-values. Instances with an adjusted *p* ≤ 0.05 were considered significant. See Supplementary Note and Supplementary Figure S10 for analysis of gene gain and gene loss in *Symbiodinium*.

## Acknowledgements

We thank Manuel Aranda for advance access to the *S. microadriaticum* genome data, and Todd LaJeunesse for information on the original isolation of *S. kawagutii*. This project was supported by the Reef Future Genomics (ReFuGe) 2020 International Consortium and the Great Barrier Reef Foundation. The data reported in this work were supported by funding from Bioplatforms Australia through the Australian Government National Collaborative Research Infrastructure Strategy (NCRIS). In memory of SF, our friend and colleague who is sorely missed.

## Author contributions

HL, MAR and CXC conceived the study and designed the experiments. HL, TGS, RGP and CXC conducted all computational analyses. VHB and BL established the algal cultures and extracted the DNA. HL, TGS, RGP, IC, MAR and CXC analysed and interpreted the results. HL and CXC prepared all figures, tables, and the first draft of this manuscript. SF and CRV provided analytical tools and scripts. HL, TGS, MAR and CXC wrote the manuscript. PB, IC, DGB, DJM, MJHvO and CRV assisted in experimental design and writing of the manuscript. All authors reviewed, commented on and approved the final manuscript.

## Data availability

All sequence data generated from this study will be available at the NCBI Short Read Archive (accessions XXXX and XXXX). Assembled genomes, predicted gene models and proteins are available at https://cloudstor.aarnet.edu.au/plus/index.php/s/6yziMf2ygWjGu0L.

## Code availability

The customised scripts for AUGUSTUS and PASA used in this study were previously published in Aranda et al.^17^; they are available at http://smic.reefgenomics.org/download/.

